# Comparison of the Accula SARS-CoV-2 Test with a Laboratory-Developed Assay for Detection of SARS-CoV-2 RNA in Clinical Nasopharyngeal Specimens

**DOI:** 10.1101/2020.05.12.092379

**Authors:** Catherine A. Hogan, Natasha Garamani, Andrew S. Lee, Jack K. Tung, Malaya K. Sahoo, ChunHong Huang, Bryan Stevens, James Zehnder, Benjamin A. Pinsky

## Abstract

**Background:** Several point-of-care (POC) molecular tests have received emergency use authorization (EUA) from the Food and Drug Administration (FDA) for diagnosis of SARS-CoV-2. The test performance characteristics of the Accula (Mesa Biotech) SARS-CoV-2 POC test need to be evaluated to inform its optimal use.

**Objectives:** The aim of this study was to assess test performance of the Accula SARS-CoV-2 test.

**Study design:** The performance of the Accula test was assessed by comparing results of 100 nasopharyngeal swab samples previously characterized by the Stanford Health Care EUA laboratory-developed test (SHC-LDT) targeting the envelope (*E*) gene. Assay concordance was assessed by overall percent agreement, positive percent agreement (PPA), negative percent agreement (NPA), and Cohen’s kappa coefficient.

**Results:** Overall percent agreement between the assays was 84.0% (95% confidence interval [CI] 75.3 to 90.6%), PPA was 68.0% (95% CI 53.3 to 80.5%) and the kappa coefficient was 0.68 (95% CI 0.54 to 0.82). Sixteen specimens detected by the SHC-LDT were not detected by the Accula test, and showed low viral load burden with a median cycle threshold value of 37.7. NPA was 100% (95% CI 94.2 to 100%).

**Conclusion:** Compared to the SHC-LDT, the Accula SARS-CoV-2 test showed excellent negative agreement. However, positive agreement was low for samples with low viral load. The false negative rate of the Accula POC test calls for a more thorough evaluation of POC test performance characteristics in clinical settings, and for confirmatory testing in individuals with moderate to high pre-test probability of SARS-CoV-2 who test negative on Accula.

## Background

The importance of diagnostic testing for severe acute respiratory syndrome coronavirus-2 (SARS-CoV-2) has been strongly emphasized by both the World Health Organization (WHO) and the United States Centers for Disease Control and Prevention (CDC) (1–3). In the US, most SARS-CoV-2 testing has been conducted using high complexity molecular-based laboratory-developed tests (LDTs) that have received emergency use authorization (EUA) by the Food and Drug Administration (FDA) in centralized laboratories certified to meet the quality standards of the Clinical Laboratory Improvement Amendments of 1988 (CLIA) (4, 5). Currently, 3 CLIA-waived point-of-care tests (POCT) are EUA-approved for SARS-CoV-2 testing: the Cepheid Xpert Xpress, the Abbott ID NOW, and the Mesa Accula (6). Compared to high complexity LDTs, POCT have the potential to reduce turnaround time of testing, optimize clinical management and increase patient satisfaction (7). The Accula SARS-CoV-2 test is a POCT that requires only 30 minutes from sample to answer and utilizes the existing palm-sized Accula dock system originally developed for rapid influenza and RSV testing. Despite the multiple potential benefits of POC assays, concern has been raised regarding their lower sensitivity for COVID-19 diagnosis compared to standard high complexity molecular based tests (8–10). It remains unclear whether this decreased sensitivity is due to test validation studies being limited to *in silico* predictions and contrived samples using reference materials, as is the case currently for the Accula SARS-CoV-2 test.

### Objectives

The aim of this study was to evaluate the test performance characteristics of the Accula SARS-CoV-2 test in a clinical setting against a high complexity reference standard.

### Study design

Nasopharyngeal (NP) swabs were collected in viral transport medium or saline from adult patients from SHC, and from pediatric and adult patients from surrounding hospitals in the Bay Area. Testing for this study was performed at the SHC Clinical Virology Laboratory using samples collected between April 7, 2020 and April 13, 2020. The same NP specimen was used for both the reference assay and Accula test for comparison.

### RT-PCR assays

The reference assay for this study was the Stanford Health Care Clinical Virology Laboratory real-time reverse transcriptase polymerase chain reaction LDT (SHC-LDT) targeting the *E* gene (11–13). The Accula SARS-CoV-2 POCT (Mesa Biotech, Inc., San Diego, CA) is a sample-to-answer nucleic acid amplification test that can yield a diagnostic result within 30 minutes of specimen collection. This test uses RT-PCR to target the nucleocapsid protein (*N*) gene, and is read out via lateral flow (**Figure 1**) (14). The manufacturer’s instructions are comprised of the following steps: collection of nasopharyngeal (NP) swab, lysis of viral particles in SARS-CoV-2 buffer, transfer of nucleic acid solution to a test cassette which contains internal process positive and negative controls, reverse transcription of viral RNA to cDNA, nucleic acid amplification, and detection by lateral flow. Due to biosafety regulations and hospital-mandated protocols for sample collection at SHC, NP swabs were directly placed into VTM or saline at the patient bedside after collection. Each test was performed at the laboratory, where a volume of 10 μL of VTM or saline was transferred to 60 μL of SARS-CoV-2 buffer and added to the test cassette. These steps were performed within a biosafety cabinet to protect against aerosolization. All remaining steps were followed as per the manufacturer’s instructions (14). Testing was repeated once for invalid results on initial testing, and the second result was interpreted as final if valid.

**Figure 1.**
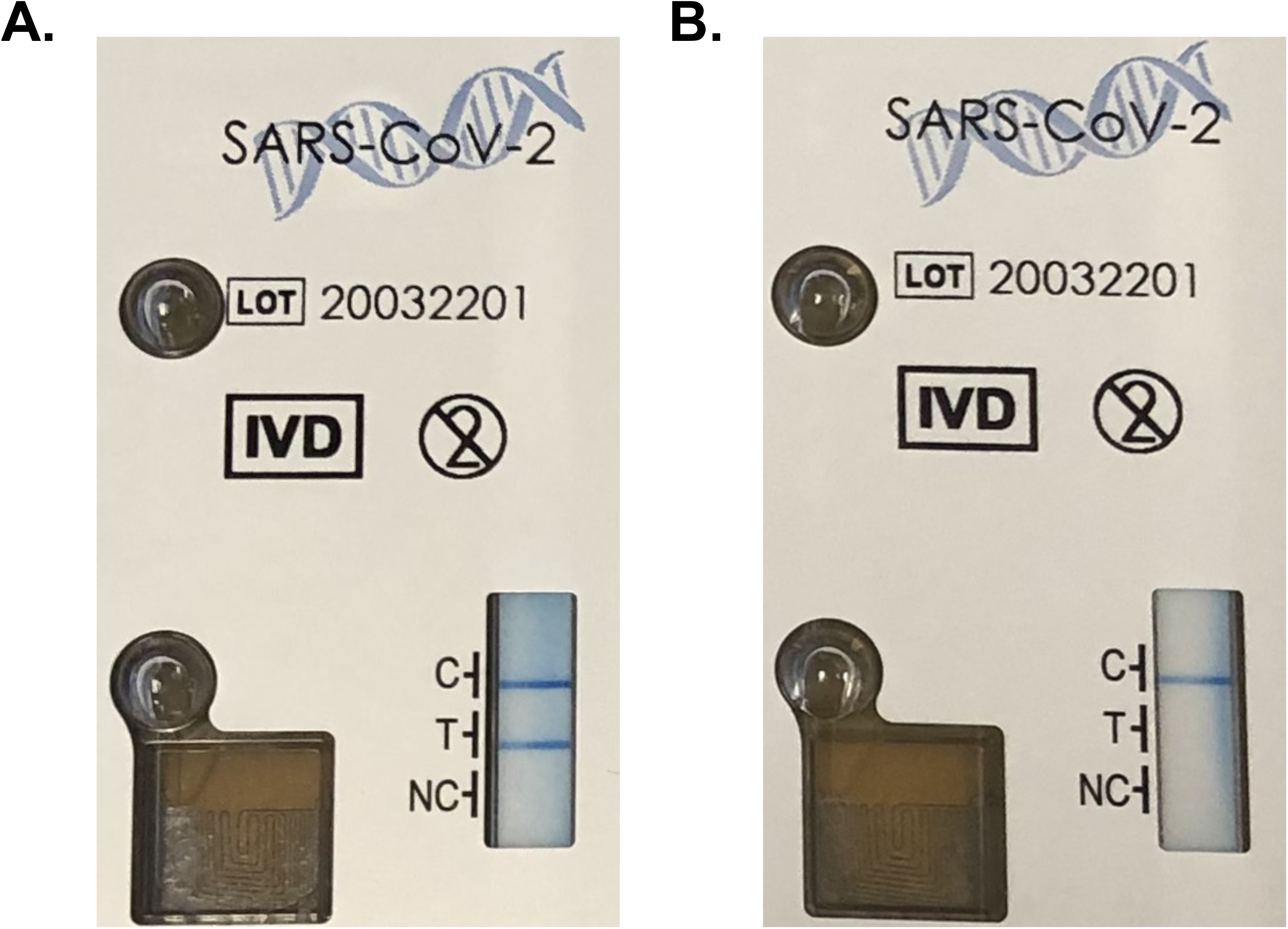
Images of the Accula SARS-CoV-2 Lateral Flow Readout. (A) positive patient specimen; (B) negative patient specimen. C, internal positive process control; T, SARS-CoV-2 test; NC, internal negative process control.

### Statistics

Overall percent agreement, positive percent agreement (PPA), negative percent agreement (NPA) and associated 95% confidence intervals (CI) were calculated. Cohen’s kappa coefficient (κ) of qualitative results (detected/non-detected) between the Accula SARS-CoV-2 test and the SHC-LDT was also calculated with 95% CI. Cohen’s kappa values between 0.60 and 0.80 were interpreted to indicate substantial agreement, and kappa calues above 0.81 were interpreted as excellent agreement (15). All analyses were performed using Stata version 15.1.

## Results

We included 100 samples (50 positive, 50 negative) previously tested by the SHC LDT, and tested in parallel with the Accula SARS-CoV-2 POCT. A total of 37 samples were collected in VTM (13 positive, 24 negative), and 63 were collected in saline (37 positive, 26 negative). Positive samples determined by the SHC-LDT included a range of cycle threshold (Ct) values, with a median Ct of 28.2 (IQR 20.4-36.3). A total of 3 samples were resulted as invalid on initial testing by Accula and were repeated once. One of these samples was detected for SARS-CoV-2 on repeat testing, and the other 2 samples were negative.

The Accula SARS-CoV-2 test correctly identified 34/50 positive samples and 50/50 negative samples, corresponding to an overall percent agreement of 84.0% (95% CI 75.3 to 90.6%), (**Table 1**). The positive percent agreement was 68.0% (95% CI 53.3 to 80.5%) and the Cohen’s kappa coefficient was 0.74 (95% CI 0.61 to 0.87), indicating substantial agreement. The 16 positive samples that were negative by the Accula test had a median Ct value of 37.7 (IQR 36.6 to 38.2) by the SHC-LDT, consistent with lower viral loads. The NPA was 100% (95% CI 92.9 to 100%). The lateral flow read-out on the Accular test was considered easy to interpret for all samples with the exception of a single known positive sample that showed a faint positive test line. Repeat testing of this sample showed the same faint test line, and was interpreted as positive.

**Table 1.** Comparison of the Stanford Health Care SARS-CoV-2 Laboratory-Developed Test and the Accula SARS-CoV-2 PCR Test

|  |  | Accula SARS-CoV-2 PCR Test |  | Total |
| --- | --- | --- | --- | --- |
|  |  | Detected | Not Detected |  |
| SHC-LDT | Detected | 34 | 16 | 50 |
|  | Not Detected | 0 | 50 | 50 |
| Total |  | 34 | 66 | 100 |

LDT: Laboratory-developed test; SARS-CoV-2: severe acute respiratory syndrome coronavirus 2; SHC: Stanford Health Care

## Discussion

Although SARS-CoV-2 testing capacity has improved in many countries, a global shortage of diagnostic infrastructure and consumable reagents has limited testing efforts. Point-of-care tests offer the potential advantages of improved access to testing and reduced turnaround time of results. Of the multiple EUA assays for diagnosis of SARS-CoV-2, only the Xpert Xpress, the ID NOW, and the Accula are CLIA-waived (6). Recent data support the test performance of the Cepheid Xpert SARS-CoV-2 assay, with agreement over 99% compared to high-complexity EUA assays (8, 16, 17). In contrast, some studies have raised concern regarding the diagnostic accuracy of the ID NOW, with positive percent agreement ranging from 75-94% compared to reference assays (8–10, 18). Given the poor diagnostic performance of the ID NOW, and uncertainty regarding availability of Xpert Xpress cartridges, the Accula system has been tauted as an interesting POCT alternative but data were previously lacking on its clinical performance. In this study, we showed that similar to ID NOW, the Accula SARS-CoV-2 test has a lower sensitivity for diagnosis of COVID-19 compared to an EUA LDT. The false negatives obtained from the Accula SARS-CoV-2 test were predominantly observed with low viral load specimens.

Given the accumulating evidence on lower diagnostic performance with 2 of the 3 CLIA-waived SARS-CoV-2 assays, it is now important to consider how best to integrate these tests in diagnostic workflows and to identify groups of individuals for whom POCT use should be prioritized. Furthermore, reagents and kits have been limited, which limits POCT capacity. Certain groups such as individuals requiring urgent pre-operative assessment including transplantation, patient-facing symptomatic healthcare workers, and individuals waiting for enrollment in a SARS-CoV-2 therapeutic trial have been identified as key groups in whom to prioritize POCT. However, for each of these scenarios and depending on the POCT used, the risk of missing a case due to low sensitivity must be considered. In individuals with moderate to high pre-test probability of SARS-CoV-2, reflex testing of negative samples on a separate EUA assay should be performed. Education of health care professionals on the limitations of SARS-CoV-2 POCT should also be implemented to ensure optimal interpretation and management of negative results.

Our study has several limitations. First, NP swabs were placed in VTM or saline at the patient bedside before loading the Accula test cassette, which may have decreased sensitivity by diluting the viral inoculum. Although this is discordant with the best recommended practice by the manufacturer, it is in line with the practice at multiple institutions with clinical laboratories that have assessed SARS-CoV-2 POCT due to biosafety concerns from risk of aerosolization (8–10, 18, 19). Second, it is possible that the use of saline instead of VTM led to poorer performance of the Accula. However, aliquots from the same sample were used for parallel testing with the EUA method, which minimizes sources of variation, and represents a pragmatic comparison given widespread VTM shortages. Finally, the lateral-flow read-out of the Accula test is generally easy to interpret; however, faint lines may be more challenging to interpret and lead to result discrepancies.

In summary, this study demonstrated that the Accula POCT lacks sensitivity compared to a reference EUA SARS-CoV-2 LDT. Careful consideration should be given to balance the potential advantages of rapid POCT to lower diagnostic accuracy. Individuals with moderate to high pre-test probability who initially test negative on the Accula test should undergo confirmatory testing with a separate EUA assay.

## Acknowledgments

We would like to thank the members of the Stanford Health Care Clinical Virology Laboratory, Department of Emergency Medicine, and Department of Medicine, Division of Infectious Disease for their hard work and dedication to patient care.

